# Energetic substrate availability regulates synchronous activity in an excitatory neural network

**DOI:** 10.1101/363671

**Authors:** David S. Tourigny, Muhammad Kaiser Abdul Karim, Rodrigo Echeveste, Mark R. N. Kotter, John S. O’Neill

**Author notes:** Corresponding authors: (DST); (MRNK); (JSO).

## Abstract

Neural networks are required to meet significant metabolic demands associated with performing sophisticated computational tasks in the brain. The necessity for efficient transmission of information imposes stringent constraints on the metabolic pathways that can be used for energy generation at the synapse, and thus low availability of energetic substrates can reduce the efficacy of synaptic function. Here we study the effects of energetic substrate availability on global neural network behavior and find that glucose alone can sustain excitatory neurotransmission required to generate high-frequency synchronous bursting that emerges in culture. In contrast, obligatory oxidative energetic substrates such as lactate and pyruvate are unable to substitute for glucose, indicating that processes involving glucose metabolism form the primary energy-generating pathways supporting coordinated network activity. Our experimental results are discussed in the context of the role that metabolism plays in supporting the performance of individual synapses, including the relative contributions from postsynaptic responses, astrocytes, and presynaptic vesicle cycling. We propose a simple computational model for our excitatory cultures that accurately captures the inability of metabolically compromised synapses to sustain synchronous bursting when extracellular glucose is depleted.

## Introduction

Accurately processing, storing, and retrieving information comes at a considerable metabolic cost to the central nervous system (1). It is currently thought that the human brain is responsible for 20% of all energy consumed by the body, whilst comprising only 2% of the total body weight (2). The amount of energy expended on different components of excitatory signaling in the brain has been estimated (3)(4), and mechanisms mediating synaptic transmission (including glutamate accumulation in vesicles) are predicted to monopolize 41% of all ATP turnover in the cortex (5). Theoretical considerations suggest that cortical networks therefore maximize the ratio of information transmitted to energy consumed (6). This finding could explain why the mean firing rate of neurons measured *in vivo* is much lower than that expected to maximize the brain’s total coding capacity, i.e. where neurons fire at approximately half their maximum rates, a behavior that is only very rarely observed in practice (3)(7). Mathematical models of energy-efficient neurotransmission led to the surprising conclusion that synaptic vesicle release probability is low and synaptic failures should occur often (8)(5). However, even with these adaptations for energetic efficiency the metabolic demands of neural networks remain a large proportion of the body’s total energy budget.

To meet this energetic demand, the cortex has evolved an extensive neurovascular coupling that can increase blood flow to regions of high activity. Astrocytes and other glial cells in contact with blood vessels are important regulators of brain energy supply and play a key role providing neurons with a readily accessible fuel source (9)(10). The nature of this neuron-astrocyte relationship remains controversial however, with conflicting theories concerning the primary substrate of resting *versus* active neural metabolism as well as the relative fluxes through metabolic pathways in the two cell types (11)(12)(13). Hemodynamic signals based on blood-oxygen-level-dependent functional magnetic resonance-based imaging (BOLD fMRI) show that oxygen uptake during neural activity is disproportionally small compared to that required for complete oxidation of glucose (i.e. 6O_2_ per glucose consumed), suggesting that glycolysis is the major metabolic pathway of active cortex (14). This observation led to a proposal that neural activity induces aerobic glycolysis in astrocytes, which then produce lactate to serve as the main fuel source for neurons (15)(16). Whilst, as an energetic substrate, lactate can support some aspects of synaptic function (17)(18), the astrocyte-neuron lactate shuttle hypothesis challenges the long-standing consensus that glucose is the principal fuel source of neuronal metabolism (12). Other recent studies continue to support the idea that significant amounts of glucose feed directly into neuronal glycolysis however (e.g. (19)(20)(21)), and advancements in fluorescent imaging demonstrate that activity stimulates glycolysis, but not lactate uptake (22).

Presynaptic nerve terminals are unusual in the sense that many lack mitochondria but are able to satisfy the sizable ATP-consumption requirements of synaptic vesicle recycling (23)(24)(25). ATP production must be able to increase rapidly in order to meet acute changes in presynaptic demands, and so it is perhaps not surprising that recent work has highlighted the importance of locally-derived glycolytic ATP generation and glucose transport in the synaptic vesicle cycle (26)(27). Neuronal activity has been associated with localization of key glycolytic enzymes to the presynaptic terminal (28), and several of these proteins are enriched in synaptic vesicles (29)(30) where they are found to be essential for synaptic vesicle re-acidification and glutamate uptake (31)(32). Vesicle recycling is a highly dynamic and energetically-demanding process (33)(34) for which ATP supply *via* glucose oxidation alone is presumably too slow (35)(36). Consequently, metabolic stress induced at individual excitatory presynapses by substrate depletion has been shown to reduce the number of functional release sites and depress rates of synaptic vesicle recovery (37)(38). It is not yet understood, however, whether these changes in central carbon metabolism directly affect global network behavior. In this study we set out to determine the consequences of energetic substrate depletion on a network model of human cortical neurons.

Our approach was to develop a combined experimental-computational model sufficiently detailed to be relevant to the problem at hand yet sufficiently simple to provide an intuitive picture of how metabolism governs important aspects of cortical network behavior. For these experiments we used induced cortical glutamatergic neurons (iNs) derived from human embryonic stem cells (hESCs) by overexpression of neurogenin 2 (NGN2) (39) from a genetically safe harbor in order to maximize induction efficiency and improve network homogeneity (40). We cultured human iNs together with rat astrocytes on multi-electrode arrays (MEAs); extensive characterization of their electrophysiology has been performed previously (39)(41) and suggests individual iNs constitute excitatory cortical layer 2/3 neurons equipped with AMPA receptors. For computational conceptualization of our experimental results we extended a simplified version of the spiking neural network model used by Guerrier *et al.* (2015) to describe the emergence of synchronous bursting driven by synaptic dynamics (42). We found a reduced version of this model captured the same effects and incorporated metabolic regulation of synaptic vesicle recovery in order to interpret experimental data derived from cultured excitatory networks.

## Results

### Emergence of synchronized bursts during excitatory network development

We cultured human iNs together with rat astrocytes on MEAs from day 3 of induction onwards. From day 12 we recorded 10min of electrical activity at three regularly-spaced intervals (10:00 AM GMT every Monday, Wednesday, and Friday) each week for a total of six weeks (Materials and Methods). To ensure the observed network behavior was representative we repeated this experiment on three separate occasions, each time using a different hESC clone for induction and a new preparation of astrocytes for co-culture. The emergence of spontaneous bursting after 3-4 weeks from induction was a consistent feature of developing networks (Fig. 1). Bursts were easily identified upon visual inspection of recorded data and consisted of a characteristic, high-frequency spike train (Fig. 1A). Timings of bursts were synchronized across all participating electrodes whilst spontaneous action potentials occurring within and between bursts were not. A custom-built synchronous burst detection algorithm (SI Appendix) was used to analyze bursts from raw electrode data and showed that synchronous bursting spread and then stabilized across the network over the course of development. Network maturation was accompanied by a gradual increase in synchronous burst frequency (Fig. 1B), but precise frequencies varied significantly between cultures. Although we do not present an extensive analysis here, we also experienced that, during later stages of network maturation, synchronous bursts underwent different degrees of higher level organization, including the appearance of compound bursting (43) and burst compactification (44).

**Figure 1.**
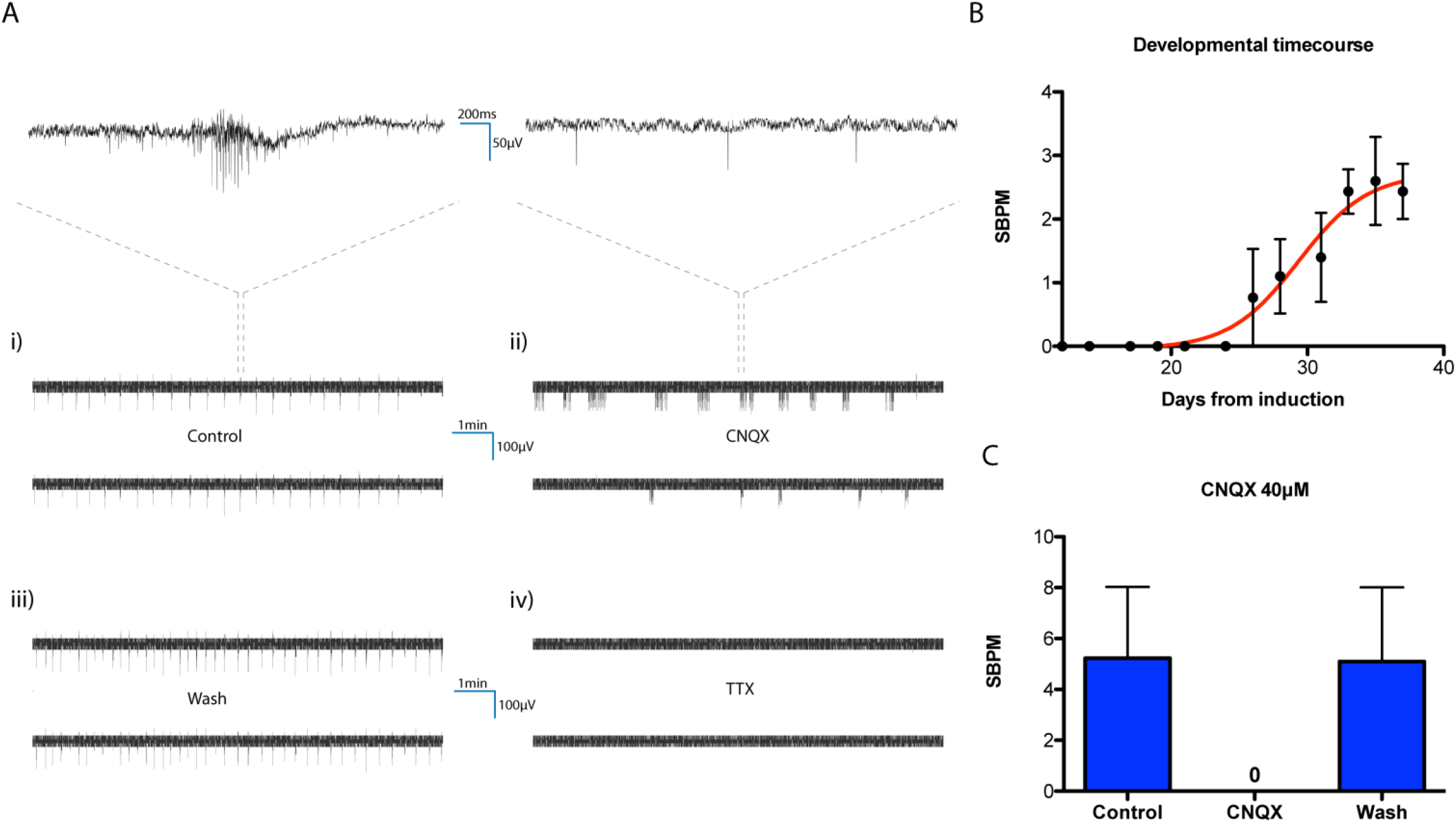
Synchronous bursting characteristics. **A)** Synchronized bursts consisting of trains of action potentials are clearly visible in raw electrode data. Representative data from two neighboring electrodes showing temporal correlation of bursts occurring in control conditions (i). Synchronized bursts are obliterated (although spontaneous action potentials persist) by the presence of the AMPA/kainite receptor antagonist CNQX (ii), but this effect is immediately reversible following a wash-off (iii). Total inhibition of electrical activity upon treatment with TTX (iv). For illustrative purposes, upper panels displaying fast time scale are smoothed using a 2ms Gaussian window. **B)** Numbers of synchronous bursts per minute (SBPM) gradually increases and stabilizes over the course of network development (data from three independent 6-week-long experiments each using distinct hESC clones and astrocyte preparations). Two-way ANOVA reports time (days from induction) as the major source of variation, P_time_ < 0.0001; N=3. **C)** When mature, the same cultures were subject to CNQX treatment, revealing the dependence of synchronous bursting on excitatory glutamatergic signaling. Zero synchronous bursts were observed in the presence of CNQX.

Several groups have similarly described the emergence of network-wide synchronous bursts in cultures of disassociated primary rat neurons (45)(46)(47) and human iNs (48). The frequency of synchronous bursting reported by Frega et al. (4.1 ± 0.1 burst/min) (48) lies within the SBPM range that we observe in mature cultures, suggesting that the two phenomena are closely related. The similar characteristics of synchronous bursting are perhaps expected given that iNs were used in those experiments also, but one cannot rule out possible differences caused by viral targeting or contributions from excitatory/inhibitory neuronal contamination in astrocyte preparations (49). To confirm that synchronous bursts are dependent on excitatory glutamatergic signaling we subjected each culture to treatment with 40μM cyanquixaline (CNQX) following the developmental time course. CNQX is a specific, competitive inhibitor of excitatory AMPA/kainite receptors (50). 10min incubation of cultures in the presence of this drug resulted in total inhibition of synchronous bursts without affecting spontaneous firing of action potentials (Fig. 1A, 1C and S1) (see also Supplementary Dataset 1). The effect was immediately reversible following a single wash with fresh media. Subsequent administration of 1μM tetrodotoxin (TTX), a potent voltage-gated sodium channel blocker, silenced the network entirely (Fig. 1A and S1) (see also Supplementary Dataset 1). Conversely, administration of 40μM bicuculline, a competitive antagonist of the primary inhibitory GABA receptor, had no detectible effect on synchronous bursts (not shown).

### Kinetics of vesicle re-acidification determine synchronous bursting frequency

Having confirmed that neuronal communication *via* release of the excitatory neurotransmitter glutamate is responsible for coordinating synchronous bursting, we were interested to know what aspects of neurotransmission determine synchronous bursting frequency. To evaluate the role of the synaptic vesicle cycle we sought to inhibit a process known to be important for vesicle maintenance and recovery following exocytosis. Various pathways and vesicle pools are suggested to participate in synaptic vesicle recycling (33)(34), and the currently accepted knowledge is concisely summarized by KEGG pathway entry hsa04721. Common to all pathways is vesicle re-acidification by the vacuolar-type ATPase (v-ATPase) required to generate the electrochemical proton gradient that is an essential prerequisite for the uptake of glutamate into synaptic vesicles (51)(52).

We evaluated the response of cultures to acute pharmacological inhibition of the v-ATPase by rounds of 10min incubation in media treated with drug or vehicle, transfer to the MEA recording device for a 200sec equilibration, and a 10min recording of network activity (Materials and Methods). We found that 50nM concanamycin A (CMA), a highly specific inhibitor of the v-ATPase (53), significantly reduced the number of synchronous bursts per minute (SBPM) (Fig. 2, vehicle: 4.76 ± 1.76; CMA: 0.30 ± 0.15) suggesting that vesicular re-acidification is a key determinant of the frequency of network-wide synchronous bursts. Conversely, treatment with 80μM dynasore (54), an inhibitor of all the major dynamin isoforms required for vesicle endocytosis, had no significant effect after the first or second 10min incubation (Fig. 2B, vehicle: 3.90 ± 1.20; dynasore 10min: 4.33 ± 1.11; dynasore 20min: 4.00 ± 1.50).

**Figure 2.**
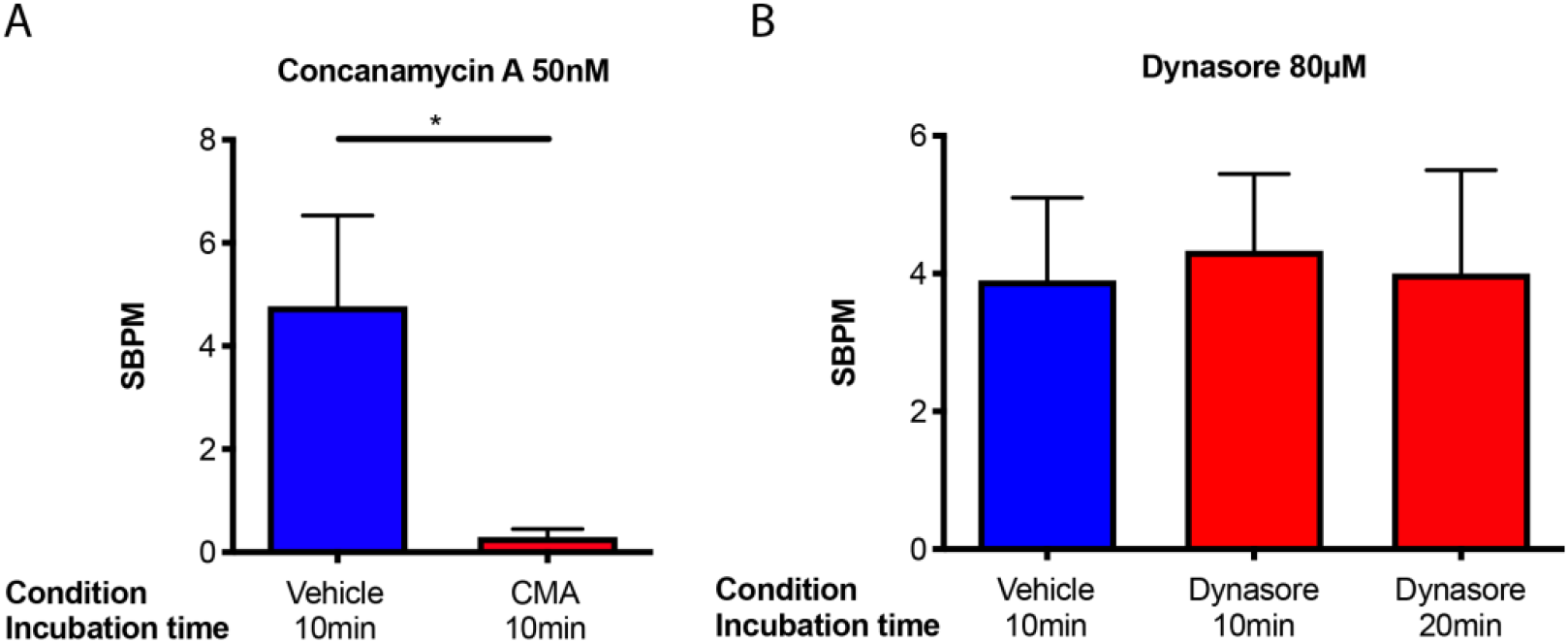
Inhibiting the vesicle recycling and maintenance pathways. **A)** Incubating cultures in the presence of 50nM CMA for 10min significantly reduces the frequency of synchronous bursting compared to 10min incubation in the presence of vehicle alone. CMA *versus* vehicle P = 0.032; N=3. **B)** 80μM dynasore had no detectible effect on SBPM after the first, or second, 10min incubation. Dynasore 10min *versus* vehicle P = 0.4, dynasore 20min *versus* vehicle P = 0.48; N=3.

### Glucose depletion reduces synchronous burst rate

To assess the consequences of energetic substrate restriction on synchronous bursting we developed a timing-based protocol of substrate depletion and repletion (Fig. 3A and Materials and Methods). On the basis of recent work, 20-30min in the absence of extracellular glucose is sufficient to impair presynaptic transmission (55)(37)(38), which is far shorter than the 16h time window during which the survival of cultured neurons remains uncompromised (56). As with previous pharmacological experiments, we repeated multiple rounds of 10min incubation, 200sec equilibration, followed by a 10min recording after which transfer of a culture into fresh media was always performed regardless of whether or not it contained an alternative mix of substrates (Fig. 3A and Materials and Methods). This was to rule out any confounding effects of mechanical perturbation or media acidification/oxygenation. We found that synchronous burst frequency (as measured by SBPM) decreased following the first 10min incubation in the absence of 25mM glucose and 0.22mM pyruvate and was significantly reduced after the second 10min incubation, with far fewer synchronous bursts occurring at regularly-spaced intervals across the subsequent 10min recording (Fig. 3B and S2, glucose and pyruvate: 6.23 ± 2.00; no substrate 10min: 4.60 ± 1.01; no substrate 20min: 0.30 ± 0.12) (see also Supplementary Dataset 2). This phenomenon was significantly reversed after the reintroduction of 25mM glucose alone (Fig. 3B and S2, glucose alone: 3.95 ± 1.20) (see also Supplementary Dataset 2), indicating that a high SBPM can be sustained in absence of the oxidative substrate pyruvate.

**Figure 3.**
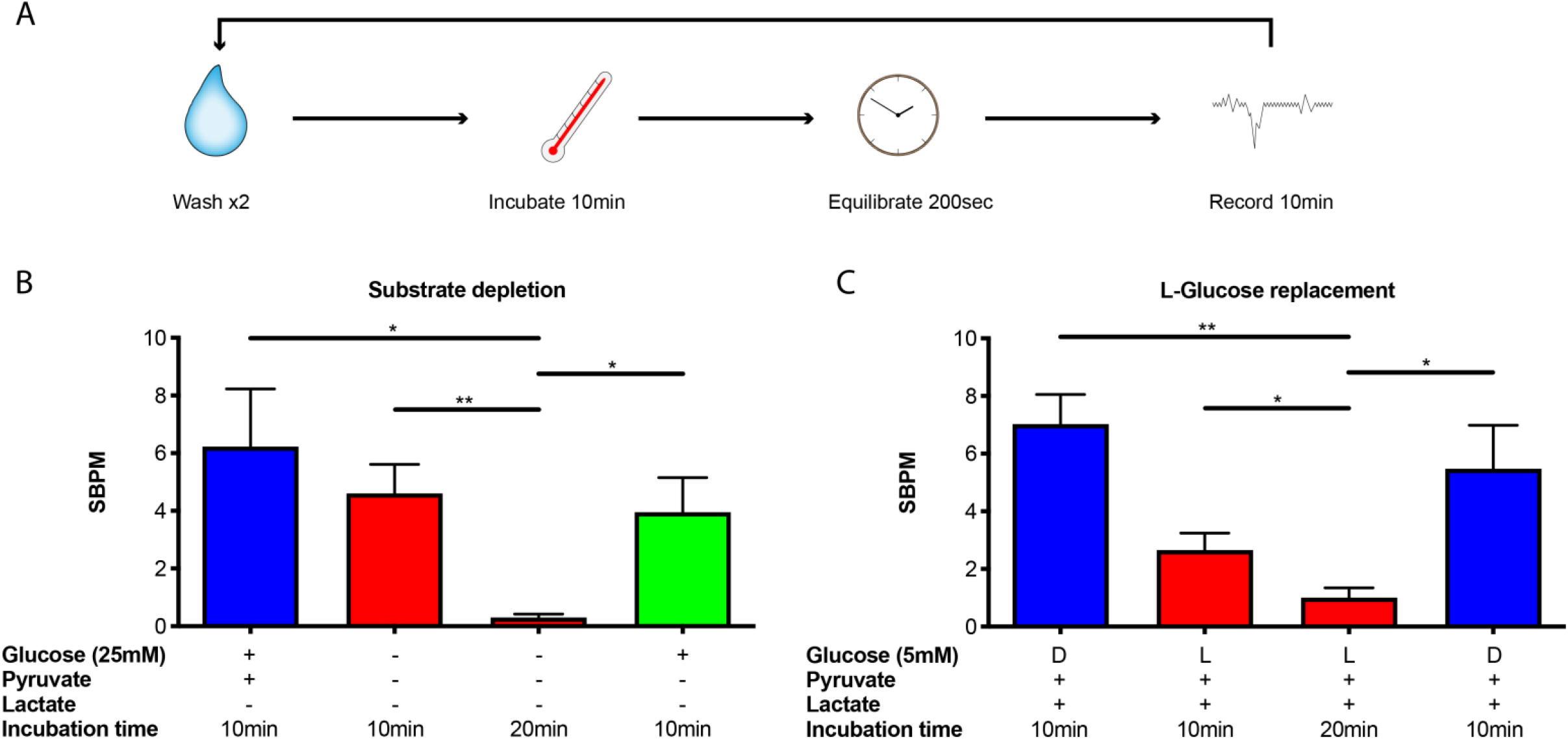
Glucose depletion decreases SBPM. **A)** Cartoon schematic of substrate depletion-repletion experimental protocol described in Materials and Methods. Each wash-incubate-equilibrate-record epoch was performed using fresh media regardless of substrate composition. **B)** Synchronous burst frequency decreases following 10min incubation in the absence of extracellular glucose (25mM) and pyruvate (0.22mM), and significantly further still following a second 10min incubation. Replenishment of glucose alone is sufficient to restore a significantly higher SBPM. No substrate 20min *versus* glucose and pyruvate P = 0.013; no substrate 20min *versus* no substrate 10min P = 0.003; glucose alone *versus* no substrate 20min P = 0.012; N=4. **C)** Only the metabolically active D-isoform of glucose (not L-glucose) can sustain a significantly higher SBPM in physiologically-relevant conditions containing 5mM D- or L-glucose, 5mM DL-lactate (racemic mixture) and 0.22mM pyruvate. L-glucose 20min *versus* D-glucose control P<0.001; L-glucose 20min *versus* L-glucose 10min P = 0.027; D-glucose replenishment *versus* L-glucose 20min P = 0.014; N=4.

In order to confirm our observations are not related to the artificially high levels of glucose (25mM) conventionally used in cell culture media, a consequence of osmotic stress possibly experienced upon exchange of the growth media, or complete absence of an alternative oxidative substrate altogether, we repeated the substrate depletion-repletion protocol under more physiologically-relevant conditions (57). We retained 0.22mM pyruvate and supplemented with 5mM DL-lactate (racemic mixture) at all time points, reducing the total concentration of extracellular glucose to 5mM. To control for the possible effect of osmotic stress during periods of glucose depletion we replaced D-glucose with its non-metabolically-active isoform L-glucose. In accordance with previous experiments, we found that substituting 5mM D-glucose with 5mM L-glucose led to a slight decrease in synchronous burst frequency after 10min incubation followed by a significant drop in SBPM during the second 10min recording (Fig. 3C and S3, D-glucose: 7.02 ± 1.02; L-glucose 10min: 2.65 ± 0.60; L-glucose 20min: 1.0 ± 0.35) (see also Supplementary Dataset 3). This effect was significantly reversed upon replacement of L-glucose with D-glucose (Fig. 3C and S3, D-glucose replenishment: 5.48 ± 1.51) (see also Supplementary Dataset 3), confirming that the metabolically active form of glucose alone can sustain a high synchronous bursting frequency. The continued presence of substrates that can only be used to generate ATP by oxidative phosphorylation implies that the glycolytic substrate glucose is required to sustain a high SBPM. Specificity for glucose was further supported by substituting 5mM glucose for 5mM galactose, a glycolytic substrate whose transport and pre-processing means it passes through glycolysis more slowly, but yields the same ATP molar equivalent to glucose during oxidative phosphorylation (Chapter 16.1.11. in (58)), which also failed to rescue the higher synchronous bursting frequency (Fig. S4) (see also Supplementary Dataset 4).

Finally, to conceptualize how these features lead to the emergence of synchronous bursting initiated by spontaneous action potential firing we have built upon the results of Guerrier *et al.* (42) to describe our excitatory network with a model based on random, sparse synaptic connections equipped with short-time synaptic plasticity (STSP), three synaptic vesicle pools, and a mechanism linking vesicle maintenance and recovery rates to energetic substrate availability. In our implementation of this computational model we explored the possibility of simplifying STSP dynamics further and found that the assumption of just a single vesicle pool (Fig. 4A) remains sufficient to recapitulate the synchronous bursting phenotype we observed experimentally during glucose depletion (Fig. 4B, Material and Methods and SI Appendix).

**Figure 4.**
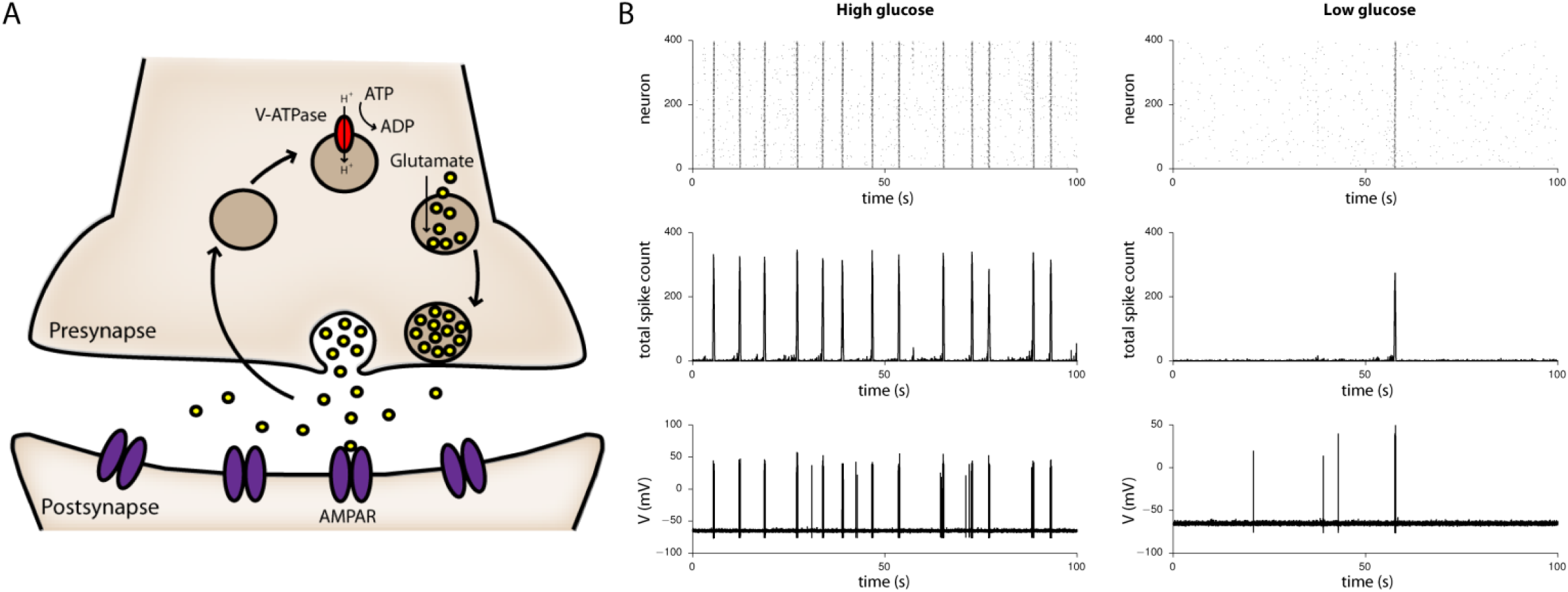
Computational model of synchronous bursting. **A)** Cartoon schematic showing that presynaptic vesicle recycling involves several steps suggested to depend on ATP supplied by glycolysis, including endocytosis and vesicle re-acidification by the v-ATPase. In our computational model we encompassed contributions from all pathways in a single term describing the rate of synaptic vesicle recovery and maintenance. **B)** Simulations reveal that in high levels of extracellular glucose (left), synchronized bursting persists at a higher frequency than when extracellular glucose is low (right), as modelled by reducing the rate of vesicle recovery and maintenance. Upper panels display raster plots of spike timings from all 400 neurons in the simulated network, middle panels the total spike count across the network as a function of time, and lower panels the corresponding fluctuations in membrane potential of a representative neuron.

## Discussion

In this work we used cultured networks of excitatory human iNs and rat astrocytes as an experimental model system to study the influence of energetic substrate availability on global network behavior. Glucose depletion dramatically reduced the prevalence of network-wide synchronous bursts mediated by excitatory neurotransmission, an effect that could not be rescued by a purely respiratory substrate such as lactate or pyruvate. Our results build upon earlier work focused on the effects of energetic substrate availability on neuronal function, which revealed that glucose is essential for synaptic transmission even though intracellular ATP levels remain normal in the presence of oxidative fuel sources (59)(60)(61)(62).

Using CNQX to inhibit synaptic transmission we found that network-wide synchronous bursts are an emergent property of excitatory synaptic communication rather than synchronization of intrinsically bursting neurons. In a simple computational model, we demonstrated that this type of synchronized bursting is sensitive the rate of synaptic vesicle recycling and maintenance, which is in turn sensitive to energetic substrate availability. Our model revealed that a single vesicle pool was sufficient to recapitulate our experimental observations, which is particularly important given the lack of conclusive experimental evidence for the involvement of different pathways and vesicle pools during synaptic vesicle recycling (33)(34), about which we prefer to remain agnostic. An exception however, is our acceptance that synaptic vesicle re-acidification is a central component of all vesicle recycling and maintenance pathways, which explains the decrease in SBPM we observed following pharmacological inhibition of the v-ATPase. Our study therefore suggests that vesicle re-acidification plays an important role as a determinant of synchronous bursting frequency.

Recent lines of evidence indicate that fully-functional presynaptic transmission is dependent on activity-induced glycolysis (31)(32)(26)(28) and this has led to a proposal that a rapid supply of ATP is required to power the synaptic vesicle cycle at nerve terminals (36). This claim is further supported by studies demonstrating a decrease in vesicle recovery rates and impaired vesicle maintenance within individual synapses in the absence of extracellular glucose (19)(37)(38), but we note other factors such as the inability of the presynaptic action potential to trigger vesicle release cannot be fully excluded (55). Our experimental results are consistent with the idea that efficacy of the synaptic vesicle cycle may diminish upon a drop in extracellular glucose concentration, and as a proof-of-concept we have accommodated this in our computational description of excitatory cultures. Following the approach of Lucas *et al.* (38), we modified the time constant for vesicle recovery in the model to account for the fact that under low glucose conditions, when energy charge (ATP:ADP/AMP ratio) is more sensitive to increased ATP consumption rates, the proportion of functional vesicles available for release is reduced (31)(32)(26)(28)(27). As observed experimentally, when vesicle recovery and maintenance is compromised in response to a decrease in extracellular glucose concentration, there is a corresponding decrease in synchronous burst frequency (Fig. 4B). A possible explanation for why a reduction in glucose availability might impair the synaptic vesicle cycle is that v-ATPase activity is more reliant on glycolytically-derived ATP (31)(32). However, other stages of vesicle recycling are likewise energetically demanding processes that depend on both oxidative (63) and non-oxidative supply of ATP (26)(28). Therefore, although it has been argued that availability of vesicles does not become rate-limiting during ATP depletion (64)(65), and despite the fact that in our hands the inhibitor dynasore had no effect on synchronous bursting frequency, the impairment of endocytosis upon glucose restriction, particularly involving rapid “kiss-and-run” (34) or dynamin independent (66) mechanisms, cannot be completely ruled out.

Dependence on glucose might be relevant for presynaptic function beyond the synaptic vesicle cycle (67) because many nerve terminals are thought to lack mitochondria (23)(24) suggesting that a considerable proportion of presynaptic ATP supply may be glycolytic in origin. Our experiments do not exclude the possibility that postsynaptic function also becomes compromised by the removal of glucose since reduced excitatory postsynaptic potential propagation also leads to a decrease in SBPM in our computational model. Postsynaptic compartments of neurons have considerable energy requirements associated with reversal of ion fluxes and membrane potential maintenance, but ATP for these processes is thought to be supplied almost exclusively *via* oxidative metabolism (3)(4)(5). The inability of pyruvate or lactate to sustain a high SBPM in our experiments suggests that the observed dependence on glucose is not limited to its role as a substrate for oxidative metabolism however, because these substrates have been shown to support many aspects of neuronal function (17)(18)(38) that can be left to depend exclusively on ATP supplied by oxidative phosphorylation. Galactose was also unable to sustain a high SBPM, implying that glycolytically-derived ATP is likely an important energy contributor to synchronous bursting. Alternatively, it has been hypothesized that astrocytically-derived lactate can be used as a substrate for oxidative metabolism by neurons (15)(16), and thus in principle could become rate-limiting for neuronal activity during glucose depletion, in a fashion that simply cannot be rescued by the presence of lactate and pyruvate in the extracellular media. As well as providing vital support for synaptic function, the presence of astrocytes in co-cultures is known to critically shape the metabolic profile of both neurons and astrocytic metabolic gene-expression profiles (68) that may in turn affect the glycolytic capacity of both cell types *in vitro*. How well these expression patterns correspond to those of intact brain is currently not completely clear however (67), and the finding that culture microenvironments potentially alter preferences in bioenergeic pathway use (69) should be taken into consideration when using experimental results to infer the relative contributions of various metabolic pathways *in vivo*. In addition, metabolic reprogramming from aerobic glycolysis to oxidative phosphorylation has been shown to occur during neuronal differentiation (70) meaning dependence on glucose as a non-oxidative fuel source may depreciate after further maturation.

It is also important to highlight the effects that glucose depletion can exert on neuronal activity through cellular signaling. Most likely this would occur indirectly *via* the AMP-activated protein kinase pathway that regulates the activity of proteins involved in fuel supply and ATP turnover in response to changes in energetic demands (71). In the brain there is no good evidence for a direct glucose-sensing mechanism such as that thought to exist in pancreatic β-cells (72), but it is understood that many neuronal cell types express ATP-sensitive potassium channels that provide an additional level of coupling between intracellular energy status and membrane excitability (73)(74). This finding has been suggested to underpin the effect that ketogenic diets can have to reduce risk propensity to epileptic seizures (75). As such, treatment with inhibitors of glycolysis including 2-deoxy-D-glucose has recently been suggested as a route towards effective seizure management (76). The rationale for such treatments is based on the observation that inhibition of glycolysis suppresses network excitability and epileptiform bursting both *in vivo* and in hippocampal slices (77)(78)(79), which complements the results we present here showing that glucose depletion decreases synchronous bursting frequency in cultured networks and that this can potentially be attributed to glycolytic cessation.

In summary, our results show how network-wide synchronous activity emerging from excitatory coupling and synaptic vesicle dynamics is regulated by energetic substrate availability in a simplified cultured network model of neurons and astrocytes. The failure to sustain a high synchronous bursting frequency in the absence of any metabolic fuel source can be explained by the fact that synaptic transmission is a highly energetically-demanding process that requires ATP for vesicle maintenance and recovery at the presynapse in addition to reversal and restoration of ion fluxes and membrane potential at the postsynapse. Sources of ATP for these processes may involve contributions from neuronal or astrocytic glycolysis, which would necessitate the particular dependence of synchronous bursting frequency on glucose. However, it is very likely that glucose oxidation also contributes toward the supply of ATP required to sustain synaptic activity. Thus, our combined experimental-computational approach paves the way for establishing an effective and pragmatic model for (dys)regulation of metabolism in the (un)healthy human brain. By making experimental data and computational code available to the wider community we hope to contribute to the further advancement of knowledge on this important subject.

## Materials and Methods

### Cell culture

Human embryonic stem cells H9 [Wisconsin International Stem Cell (WISC) Bank, WiCell Research Institute, WA09 cells] were cultured according to WiCell stem cell protocols in 6-well plates on Matrigel (Corning, hESC-Qualified) in StemFlex (Gibco). Use of stem cell line H9 was approved by the Steering Committee of the UK Stem Cell Bank and for the Use of Stem Cell Lines (ref: SCSC18-05).

Primary mixed glial cultures were derived from P0-P2 neonatal Spraque Dawley rats and were generated along the previous guidelines (80), with minor modifications (81). The pups were euthanized following Schedule 1 rules and regulations from the Home Office Animal Procedures Committee UK (APC). Mixed glia cells were maintained for 10 days in culture after which flasks were shaken for 1h at 260rpm on an orbital shaker to remove the loosely attached microglia, and then overnight at 260rpm to dislodge oligodendrocyte precursors. Astrocyte cultures were then maintained in glial culture medium (Dulbecco’s modified eagle’s medium (DMEM) supplemented with 10% fetal bovine serum (FBS), glutamine and 1% pen/strep) and passaged at a ratio of 1:3, every 10-14 days. Cells were passaged at least once before co-culturing with iNs and were only used between passages 2 and 5.

### Induction of NGN2- hESCs and culturing iNs on MEAs

Gene targeting and generation of dual GSH-targeted NGN2 OPTi-OX hESCs was performed as described previously (40). The day prior to initiation of the reprogramming process (day 0), NGN2-hESC colonies grown to 70-80% confluency were disassociated using Accutase (Sigma-Aldrich) and isolated NGN2- hESCs were seeded in 6-well plates at a density of 25,000cells/cm^2^ on Matrigel (Corning, hESC-Qualified) in StemFlex (Gibco) supplemented with RevitaCell (Gibco). Inducible overexpression of NGN2 began on day 1 by transferring cells into D0-2 induction medium (DMEM/F12, 1% Pen/Strep, 1x non-essential amino acid solution (NEAA, Gibco), 1% (v/v) N-2 supplement (Gibco), 1x Glutamax (Gibco), and doxycycline (dox) at 4μg/ml). On day 3, cells were dissociated using Accutase and re-suspended at a density of 4000cells/μl in >D2 medium (Neurobasal-A Medium (ThermoFisher), 1% pen/strep, 1x Glutamax, 1x B27 supplement (Gibco), 10ng/ml brain-derived neurotrophic factor (BDNF), 10ng/ml human recombinant neurotrophin-3 (NT-3), and dox at 4μg/ml) supplemented with RevitaCell. Rat astrocytes were dissociated using 0.05% trypsin-ethylenediaminetetraacetic acid (EDTA) and re-suspended at a density of 4000cells/μl in >D2 medium supplemented with RevitaCell. Astrocytes were then mixed at a ratio of 1:1 with dox-treated NGN2- hESCs in >D2 to make a final seeding density of 2000cells/μl for each cell type.

Cells were seeded onto a selection of 8×8 (60 electrodes in total, excluding corners) electrode MEAs (60MEA200/30iR-Ti or 60ThinMEA100/10iR-ITO, Multichannel Systems) covered by Teflon-sealed lids (Multichannel Systems) and pre-coated overnight at 4°C with 500μg/ml poly-D-lysine (PDL) in ultrapure water. For seeding, MEAs were first incubated for 1h at 37°C, 5% CO_2_ with a 20μl drop of laminin solution (20μg/ml laminin in DMEM) covering the electrode region. The laminin drop was aspirated immediately prior to seeding and replaced with a 15μl drop of re-suspended cell mixture (total density of 4000cells/μl). MEAs were incubated with 15μl drops of cell suspension for 1h at 37°C, 5% CO_2_ to allow cells to adhere before being topped up to 1ml with >D2 medium (without RevitaCell). Cultures were maintained throughout lifespan at 37°C, 5% CO_2_ with 500μl media replenished every second day. 2μM cytosine β-D-arabinofuranoside was added to cultures on day 5 to inhibit astrocyte proliferation and kill undifferentiated NGN2- hESCs and dox was excluded from >D2 media from day 8 onward.

### Recording procedures

For recordings, MEAs incubated at 37°C, 5% CO_2_ were transferred to the MEA recording device (MEA2100-2×60-System, Multichannel Systems). All recordings were performed in atmospheric conditions with stage and custom-built heated lid held at 37°C. Developmental time course recordings took place every second or third day, 10min after a 37°C, 5% CO_2_ incubation following half-media change. MEAs were allowed to equilibrate for 200sec on the MEA recording device and recording sessions lasted 10min with local field potentials from all electrodes sampled at 25kHz.

A strict regime of media exchange, incubation, equilibration, and recording was enforced for all pharmacological and substrate depletion-repletion experiments. For drug treatments, MEA cultures were first washed (2x total media exchange) in >D2 media with vehicle (all drugs were diluted in DMSO or ddH2O as required for treatment and corresponding concentration of the dissolving agent used as vehicle controls) to control for the effects of mechanical perturbation, and subsequently incubated at 37°C, 5% CO_2_ for precisely 10min. MEAs were then transferred to the recording device (pre-heated to 37°C) and equilibrated for 200sec prior to a 10min recording sampling at 25kHz. This procedure was repeated, immediately following each recording, under the multiple test conditions (inclusion of pharmacological compound in fresh >D2 media) necessary for each experiment. Cultures were exchanged into fresh >D2 media on termination of the final recording and returned to incubation at 37°C, 5% CO_2_. The same protocol was employed for substrate depletion-repletion experiments using >D2 media based on Neurobasal-A medium lacking glucose and sodium pyruvate (ThermoFisher). In this case media supplemented with the appropriate combinations of 25 or 5mM D- or L-glucose, 0.22mM sodium pyruvate, 5mM sodium DL-lactate (racemic mixture), and 5mM D-galactose served as test conditions.

Recorded data were processed and analyzed using the MC Rack/MC Tools software (Multichannel Systems) and a custom-built synchronous burst detection algorithm described in the SI Appendix. The algorithm calculates the number of synchronous bursts per minute (SBPM) for each recording and was used to extract this value from all biological replicates and conditions for further statistical analysis. Two-way analysis of variance (ANOVA) was performed in GraphPad Prism to assess the influence of biological conditions (time, energy substrate composition, or drug) and biological replicates (N = number of cultures) on SBPM; resulting p-values for conditions and number of replicates for each experiment are displayed in the figure legends. Student’s t-tests were used to compare mean SBPMs from pairs of biological conditions relevant to experimental interpretation; corresponding p-values are displayed in figure legends and annotated in figures with * signifying P < 0.05 and ** signifying P < 0.01. Error bars are standard error of the mean (SEM) and SBPM values reported in main text are mean ± SEM. Representational experimental data (Supplementary Datasets 1-4) have been deposited in Figshare (see Data Deposition for link and DOIs).

### Computational modelling

We simulated a modified version of the excitatory neural network model described by Guerrier *et al.* (42) consisting of 20×20 (400) connected neurons organized on a square lattice. Membrane potential of each neuron was modelled using the simplified Hodgkin-Huxley model and neurons were connected randomly according to a probability distribution that decays as a function of distance between pairs on neurons (see SI Appendix and (42)). We believe this type of random synaptic connectivity accurately reflects that which emerges in our experimental cultures over the course of development. In the original work (42), the fraction of available *free, docked*, and *recovering* synaptic vesicles are simulated, corresponding to the proposed existence of multiple vesicle pools and recovery pathways (33)(34). Here we simulated only the fraction of docked vesicles, assuming that recovery is described by a single rate constant encompassing these mechanisms and have shown that this is sufficient to recapitulate the emergence of synchronous bursting across the simulated network (SI Appendix). To model the effect of glucose depletion on synaptic vesicle recovery, we allow for modulation of this rate by energy substrate availability, assuming that when the levels of extracellular glucose are low the overall rate of recovery decreases as a function of recent presynaptic energy consumption. Although we take inspiration from a similar approach used in (38), we do not intend to model the exact functional form of this glucose dependence and instead sought only to capture the desired properties using a simplified model of presynaptic metabolism (SI Appendix).

## Acknowledgements

We thank past and present members of the O’Neill and Kotter labs for assistance and comments. We are grateful to Simon Laughlin for inspiring discussions and suggestions. DST is a Simons Foundation Fellow of the Life Sciences Research Foundation. MKAK is supported by a Daya Diri-Cambridge Trust Scholarship. Research in MRNK’s laboratory is funded by the UK Multiple Sclerosis Society and supported by a core support grant from the Wellcome Trust and Medical Research Council to the Wellcome Trust-MRC Cambridge Stem Cell Institute. MRNK holds a National Institute for Health Research Clinician Scientist Award CS-2015-15-023 (Disclaimer: This report is independent research arising from Clinician Scientist Award, CS-2015-15-023, supported by the National Institute for Health Research. The views expressed in this publication are those of the authors and not necessarily those of the National Health Service, National Institute for Health Research or the Department of Health). JSO is supported by the Medical Research Council (MC_UP_120/4).

## Author Contributions

DST conceived study and designed experiments with support from MRNK and JSO. DST and MKAK performed cell culture and developmental time course experiments; DST and MKAK performed substrate replacement and drug treatment experiments; DST analyzed the data. RE and DST developed computational model; RE performed simulations. DST wrote the paper with input from all authors.

## Data Deposition

Representational multi-electrode array data have been deposited in Figshare (https://figshare.com) and are publicly available at the following DOIs: 10.6084/m9.figshare.6741464 (Supplementary Dataset 1); 10.6084/m9.figshare.6741479 (Supplementary Dataset 2); 10.6084/m9.figshare.6741485 (Supplementary Dataset 3); 10.6084/m9.figshare.6741494 (Supplementary Dataset 4). Computational code available in GitHub: https://github.com/r-echeveste/meta_reg_net.

