## Supporting Information for "Energetic substrate availability regulates synchronous activity in an excitatory neural network"

### SI Appendix

#### 1 Synchronous burst detection algorithm

First stages of burst detection are performed post-recording using the software MC Rack (Multichannel Systems). Spikes are detected from local field potentials at each electrode using a threshold-based detector as downward excursions beyond the estimated standard deviation (calculated across a time interval of 500ms). Individual bursts are then recorded for each electrode using the MaxInterval Method [1] implemented in MC Rack with parameters *Max. interval to start burst*: 50ms; *Max. interval to end burst*: 50ms; *Min. interval between bursts*: 100ms; *Min. duration of burst*: 10ms; *Min. number of spikes in burst*: 10. The MC Rack implementation of the MaxInterval Method is as follows:

1. Scan the spike train until an interspike interval is found that is less than or equal to *Max. interval to start burst*

2. While the interspike intervals are less than *Max. interval to end burst*, they are included in the burst

3. If the interspike interval is more than *Max. interval to end burst*, the burst ends

4. Merge all the bursts that are less than *Min. interval between bursts* apart

5. Remove the bursts that have duration less than *Min. duration of burst* or have fewer spikes than *Min. number of spikes*

Individual bursts are recorded per electrode as a list of individual burst timestamps and lengths, and the MC Rack output file is converted to ASCII format using the software MC Tools (Multichannel Systems).

The ASCII file output from MC Tools is used as an input for a custom-built synchronous burst detection algorithm written in the *R* programming language. Electrodes are pre-processed to detect outliers, which are identified and excluded from further analysis if they have a number of recorded individual bursts falling more than 1.5 times the interquartile range above the third quartile or below the first quartile, or zero. Time is binned into 100ms intervals and a logical vector spanning the length of the entire recording is initialized for each remaining electrode, containing value TRUE if an individual burst is recorded for that electrode during the given time interval and FALSE otherwise. An integer-valued burst profile vector, also spanning the length of the recording, is then constructed by summing the total number of electrodes individually bursting at each time interval. Run-length encoding of consecutive intervals within the burst profile vector where four or more electrodes are individually bursting simultaneously is used to mark the start and end of synchronous bursts, which at their peak typically involve 35-45 electrodes. Typical synchronous bursts last 600-800ms whilst individual bursts at electrodes usually have a length of 80-500ms. The total count of synchronous bursts over the course of a recording is divided by length of that recording (in mins) to calculate SBPM for further analysis. SBPM is capped at a maximum of 10 to avoid over-counting in hyper-active cultures.

### 2 Computational model

#### 2.1 Network and neural model

We employ the same models for the neural network and the neural dynamics as in the work of [2], which we recapitulate in this section, but modify the model of synaptic dynamics as described in the following section. As in the original paper, we simulate a network of 400 neurons, organized on a physical grid of 20x20, where the probability of connection is a decaying function of distance. The connectivity between neurons is defined by the binary connectivity matrix $W$, where $W(i,i) = 0$,

$$\mathcal{P}\left(W_{ij} = 1\right) = exp\left[-d(i,j)^2/2\sigma^2\right], \quad \forall i \neq j, \tag{S1}$$

$d(i,j)$ returns the distance between the two units, and $\sigma$ represents the connectivity decay length.

The neural model consists of a simplified Hodgkin-Huxley model described by:

$$C\dot{V} = I_{app} - I_{Na} - I_K - I_L + I_{syn} + \sigma_\xi \dot{\xi}, \tag{S2}$$

where $V$ is the neuron's membrane potential, $C$ is the membrane capacitance, and $I_{app}$, $I_{Na}$, $I_K$, $I_L$, $I_{syn}$ represent the external, sodium, potassium, leak, and total synaptic currents, respectively. $\xi$ denotes a Wiener process, and $\sigma_\xi$ represents the standard deviation of the noise. In this simplified version of the model, only the dynamics of the potassium activation variable $n$ are computed via:

$$\dot{n} = \alpha_n(V)\,(1-n) - \beta_n(V)\,n, \quad \alpha_n(V) = \frac{1}{10\tau_n}\frac{\theta_n - V}{exp\left(\frac{\theta_n - V}{\tau_n}\right) - 1}, \quad \beta_n(V) = \eta_n\, exp\left(-\frac{V+65}{\sigma_n}\right), \tag{S3}$$

while the sodium activation variable $m$ is assumed to instantaneously adapt to:

$$m_\infty(V) = \frac{\alpha_m(V)}{\alpha_m(V) + \beta_m(V)}, \quad \alpha_m(V) = \frac{1}{\tau_m}\frac{\theta_m - V}{exp\left(\frac{\theta_m - V}{\tau_m}\right) - 1}, \quad \beta_m(V) = \eta_m\, exp\left(-\frac{V+65}{\sigma_m}\right), \tag{S4}$$

The currents, are then calculated via:

$$I_{Na} = g_{Na} m_{\infty}^{3} (0.089 - 1.1n)(V - E_{Na})$$
$$I_K = g_K n^4 (V - E_K)$$
$$I_L = g_L \; (V - E_L)$$

All parameter are presented in Table S1. No external current is applied, but frozen noise (constant in time but variable across units) of zero mean is provided to simulate heterogeneity in the neurons resting potentials ($-65 \pm 0.2mV$). The calculation of the total synaptic current is presented after the introduction of the synaptic dynamics.

#### 2.2 Metabolic-dependent synaptic dynamics

As in [2], we consider synapses endowed with synaptic facilitation, captured by facilitation variable $x$, whose dynamics evolve according to:

$$\dot{x} = \frac{X - x}{\tau_f} + k\,(1 - x) H\,(V - T), \tag{S5}$$

where $x$ increases after every action potential of the presynaptic neuron, when $V$ is above the threshold $T$ in the Heaviside function $H$, and decays to resting value $X$ with time constant $\tau_f$. Beyond the use as a facilitation variable, functionally, $x$ is generically employed in the model as a proxy for the level of recent neural activity, which becomes useful to evaluate both refractoriness and metabolic consumption, as we later describe.

In the original work of [2], three synaptic vesicle pools are simulated, representing the fraction of available *free*, *docked*, and *recovering* vesicles. Here we simulate only the fraction of docked vesicles, assuming that their recovery is simply modulated by the ratio of metabolic flux to neural activity. We have shown that, in absence of this metabolic coupling, by setting the other vesicle pools to a constant in the original model, the bursting behaviour remains unchanged, showing that the simulation of the dynamics of these two additional pools is not relevant for the purposes of the present work, and justifying this simplification. The docked vesicle pool (represented by $y_{dock}$), evolves according to:

$$\dot{y}_{dock} = \frac{1}{\tau_{dock}} \left(y_{dock}^{max} - y_{dock}\right) \mathcal{M}(x) \left[1 + \frac{x - X}{X} H\,(V - T)\right] - \frac{1}{\tau_{rel}} y_{dock} \frac{x - X}{X} H(V - T), \tag{S6}$$

where

$$\mathcal{M}(x) = y_{ref}\, exp\left(-x/m_{\phi}\right) \tag{S7}$$

represents the metabolic modulation function. We note here that we do not intend to capture the exact functional form of this coupling in nature (which is currently unknown), but simply to present a simplified model with the desired properties: the exponential ensures the sign of the function remains positive, becoming smaller as the ratio of recent past energy consumption to resource availability decreases. This ratio is here expressed as the proxy for recent neural activity $x$ over the metabolic flux $m_{\phi}$, which we assume to be a function of glucose availability. High recent activity means here a larger resource depletion, and reduced capacity for $y_{dock}$ to recover, while a higher metabolic flux results in an increased capacity for vesicle recovery.

Finally, as in [2] we consider synapses to become refractory after persistent firing of the presynaptic neuron, when $y_{dock}$ reaches a minimal depletion variable $y_{dock}^{min}$, and stays refractory until this value has recovered above this minimal value and $x$ has has also reached a reference value $x_{ref}$. We note that the equations for $x$ and for the synaptic pools in this work and in the original model are only expressed in terms of global properties of the presynaptic neuron. All synapses of the same presynaptic neuron follow the same dynamics and can be simulated as one. In particular, all synapses corresponding to the same presynaptic neuron will become refractory at the same time. Refractory synapses do not contribute to the total synaptic current each postsynaptic neuron receives. If a spike arrives at time $t_0$ to a non-refractory synapse $j$, it produces a current given by:

$$i_j(t) = K_I\,(y_{dock}(t) - y_{dock}(t_0))\;H\,(V(t) - T)\,H\left(y_{dock}(t) - y_{dock}^{min}\right) \tag{S8}$$

which integrates the number of depleted docked vesicles and includes a proportionality constant $K_I$. The two Heaviside functions ensure that currents are only produced during spikes and only if the fraction of docked vesicles has not dropped below a minimal value. The total current is then computed as:

$$I_{syn}(t) = \sum_{j\ (non-ref)} i_j(t - t_{del}), \tag{S9}$$

where a delay time $t_{del} = 1ms$ is employed.

#### 2.3 Simulations

Simulations were performed in python, employing an adaptation of the original matlab code of [2]. The full code for the simulations here presented can be found at `https://github.com/r-echeveste/meta_reg_net`.

Network dynamics were simulated for 100s, after an initial 5s transient. The evolution of the system was computed using a fourth-order Runge-Kutta method, with $dt = 0.05$ms. Simulations were conducted in two different metabolic flux conditions: a high metabolic flux case, corresponding to high levels of extracellular glucose, and a low metabolic flux case, corresponding to low levels of extracellular glucose. The parameters for both simulations are found in Table S1.

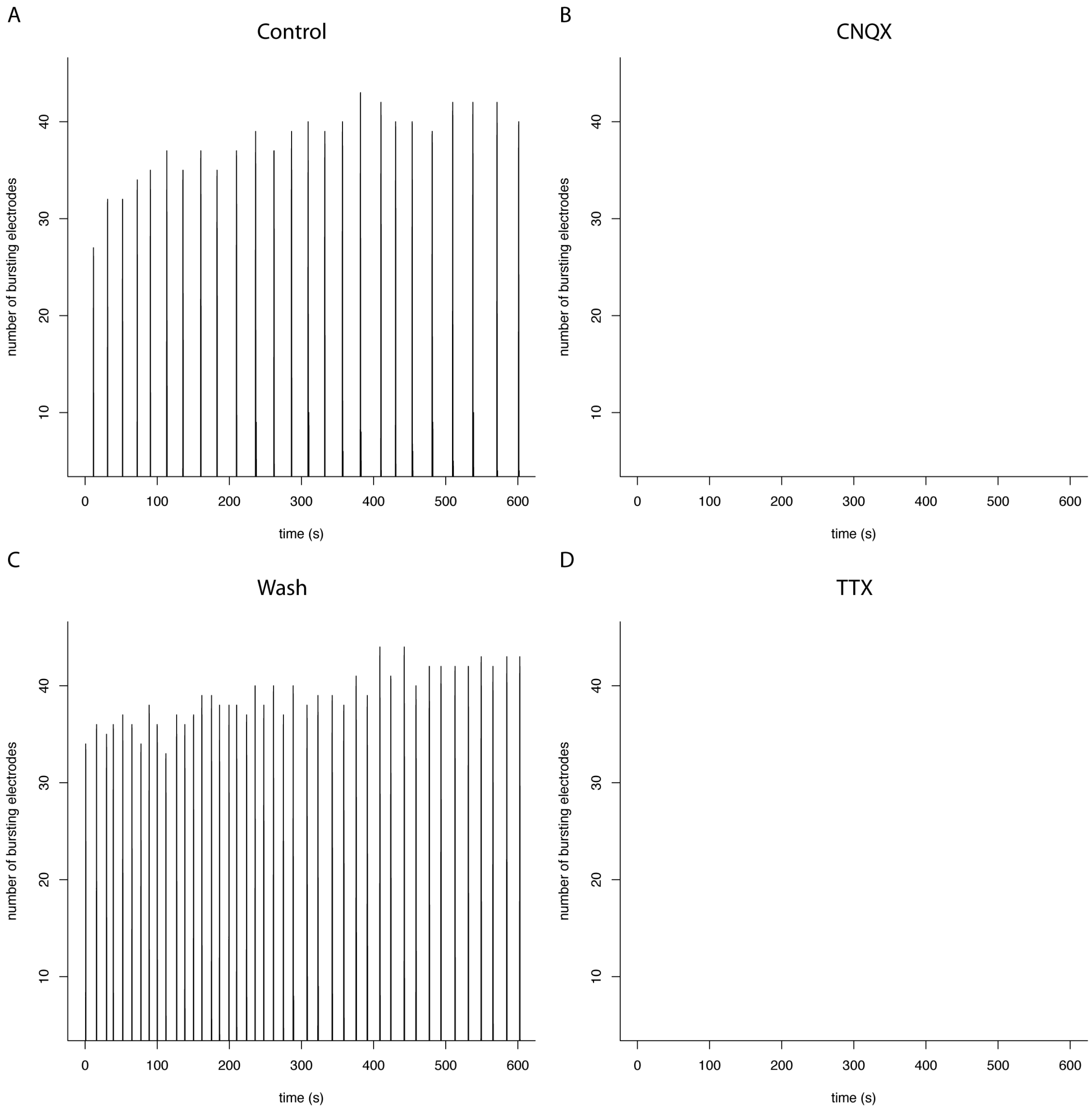


**Figure S1. Representational experimental data (pharmacological characterization).** Representational examples of the recorded data used to generate Fig. 1A in main text. These are intended to complement Supplementary Dataset 1 (10.6084/m9.figshare.6741464, each panel has an associated 600sec MEA recording). The plots represent the number of electrodes bursting across the MEA at the given time point under the given experimental conditions: **A)** control; **B)** CNQX; **C)** wash; and **D)** TTX.

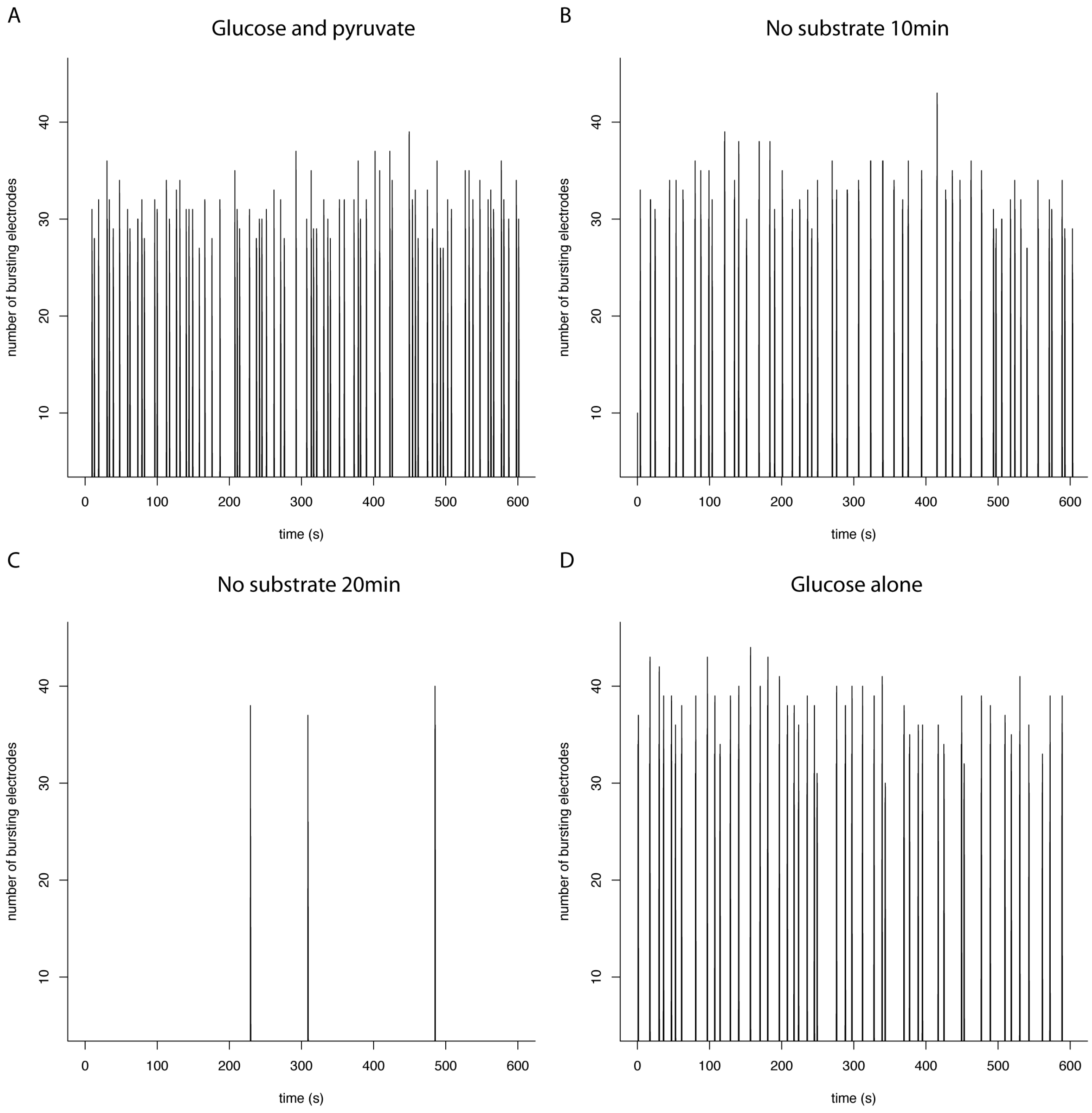


**Figure S2. Representational experimental data (substrate depletion)**. Representational examples of the recorded data used to generate Fig. 3B in main text. These are intended to complement Supplementary Dataset 2 (10.6084/m9.figshare.6741479, each panel has an associated 600sec MEA recording). The plots represent the number of electrodes bursting across the MEA at the given time point under the given experimental conditions: **A)** glucose and pyruvate; **B)** no substrate 10min; **C)** no substrate 20min; and **D)** glucose alone.

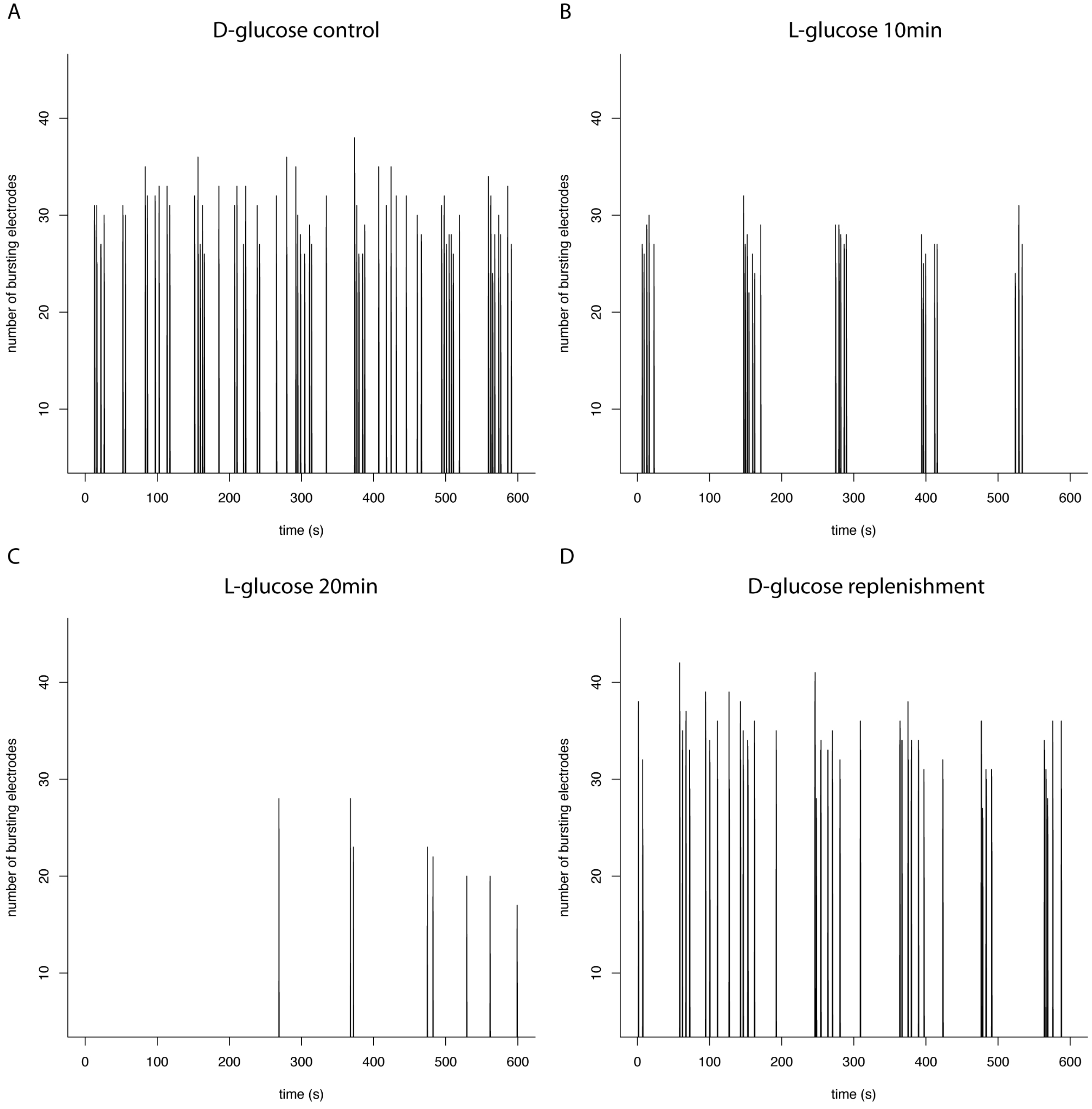


**Figure S3. Representational experimental data (L-glucose replacement)**. Representational examples of the recorded data used to generate Fig. 3C in main text. These are intended to complement Supplementary Dataset 3 (10.6084/m9.figshare.6741485, each panel has an associated 600sec MEA recording). The plots represent the number of electrodes bursting across the MEA at the given time point under the given experimental conditions: **A)** D-glucose control; **B)** L-glucose 10min; **C)** L-glucose 20min; and **D)** D-glucose replenishment.

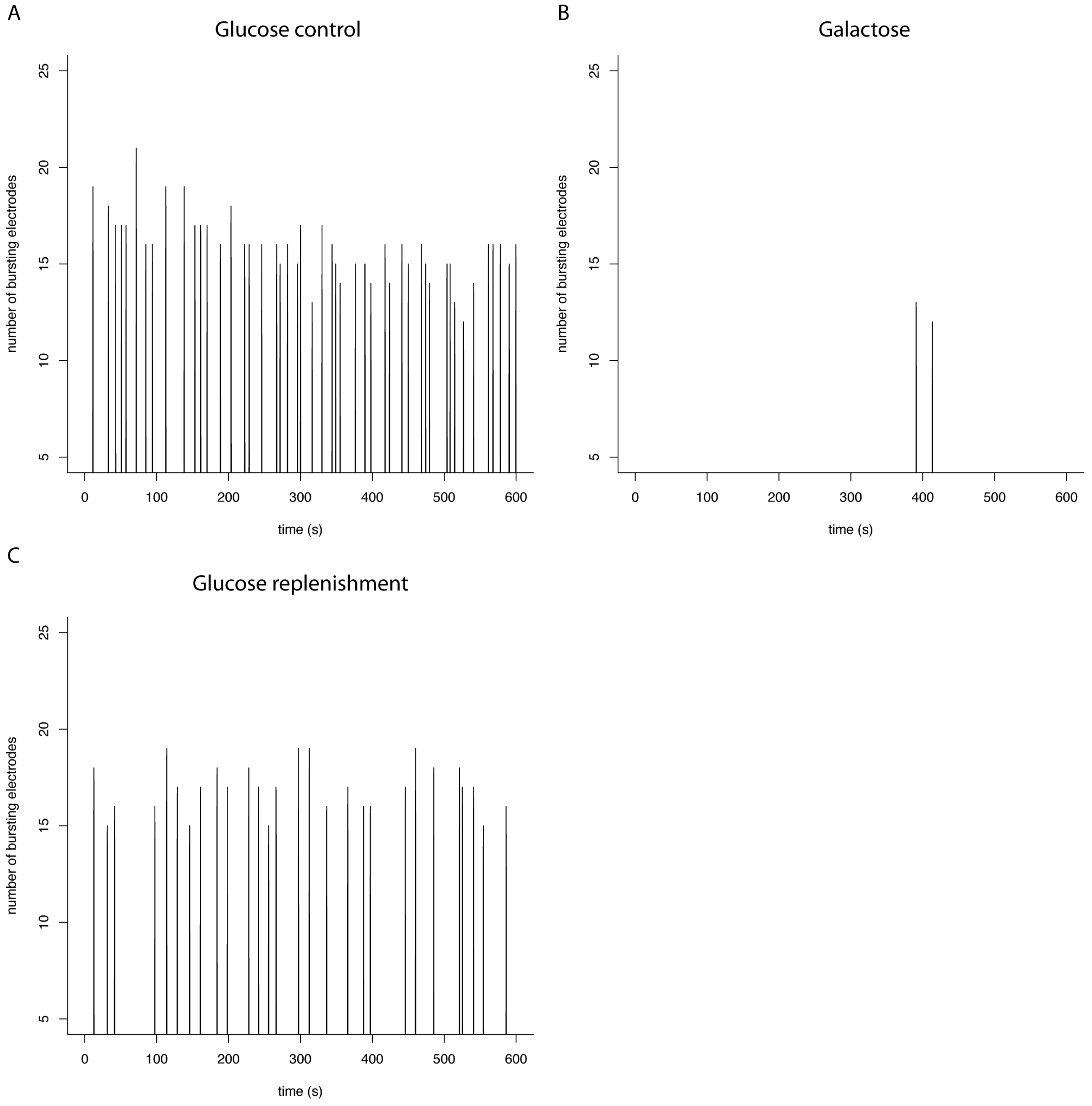


**Figure S4. Representational experimental data (galactose replacement).** Representational examples of recorded data used to study the effects of replacing 5mM glucose with 5mM galactose on synchronous bursting. These are intended to complement Supplementary Dataset 4 (10.6084/m9.figshare.6741494, each panel has an associated 600sec MEA recording). The plots represent the number of electrodes bursting across the MEA at the given time point under the given experimental conditions: **A)** glucose control; **B)** galactose; and **C)** glucose replenishment.

| Parameter | Description | Value |
|---|---|---|
| | Network | |
| $N$ | Number of neurons | 400 |
| $\sigma$ | Connectivity scale | 0.9 |
| $W$ | Connectivity matrix | random binary |
| | Neural model | |
| $C$ | Membrane capacitance | $1\mu$F cm$^{-2}$ |
| $g_{Na}$ | Conductance of Na$^{2+}$ current | 120mS cm$^{-2}$ |
| $g_K$ | Conductance of K$^+$ current | 36mS cm$^{-2}$ |
| $g_L$ | Conductance of the leak current | 0.3mS cm$^{-2}$ |
| $E_{Na}$ | Equilibrium potential of Na$^{2+}$ current | 50mV |
| $E_K$ | Equilibrium potential of K$^+$ current | -77mV |
| $E_L$ | Equilibrium potential of the leak current | -54.4mV |
| $\tau_m$ | Parameter of the m activation variable | 10ms |
| $\tau_n$ | Parameter of the n activation variable | 10ms |
| $\theta_m$ | Parameter of the m activation variable | -40mV |
| $\theta_n$ | Parameter of the n activation variable | -55mV |
| $\eta_m$ | Parameter of the m activation variable | 4 |
| $\eta_n$ | Parameter of the n activation variable | 0.125 |
| $\sigma_m$ | Parameter of the m activation variable | 18 |
| $\sigma_n$ | Parameter of the n activation variable | 80 |
| $\sigma_\xi$ | Membrane noise size | $0.89\mu$A ms$^{-1/2}$ cm$^{-2}$ |
| $T$ | Spiking threshold | -58mV |
| | Synaptic dynamics | |
| $X$ | Facilitation equilibrium value | 0.3 |
| $\tau_f$ | Facilitation time constant | 700ms |
| $k$ | Facilitation increase constant | 0.08 |
| $x_{ref}$ | Refractoriness threshold | 0.31 |
| $y_{dock}^{max}$ | Max. docked pool size | 0.18 |
| $y_{dock}^{min}$ | Min. docked pool size | 0.04 |
| $\tau_{dock}$ | Docking time constant | 738ms |
| $\tau_{rel}$ | Release time constant | 47ms |
| $K_I$ | Synaptic strength | 2666 |
| $t_{del}$ | Synaptic delay | 1ms |
| | Metabolic coupling | |
| $y_{ref}$ | Recovery reference value | 0.32 |
| $m_\phi$ | Metabolic flux (high glucose) | 1.0 |
| $m_\phi$ | Metabolic flux (low glucose) | 0.1 |
| | Simulation | |
| $dt$ | Time step | 0.05ms |

**Table S1.** List of parameter values.
